# Single-cell transcriptome analysis identifies distinct cell types and intercellular niche signaling in a primary gastric organoid model

**DOI:** 10.1101/190132

**Authors:** Jiamin Chen, Billy T Lau, Noemi Andor, Sue M Grimes, Christine Handy, Christina Wood-Bouwens, Hanlee P Ji

**Author notes:** To whom correspondence should be addressed. Hanlee P. Ji.

## Abstract

The diverse cellular milieu of the gastric tissue microenvironment plays a critical role in normal tissue homeostasis and tumor development. However, few cell culture model can recapitulate the tissue microenvironment and intercellular signaling *in vitro*. Here we applied an air-liquid interface method to culture primary gastric organoids that contains epithelium with endogenous stroma. To characterize the microenvironment and intercellular signaling in this model, we analyzed the transcriptomes of over 5,000 individual cells from primary gastric organoids cultured at different time points. We identified epithelial cells, fibroblasts and macrophages at the early stage of organoid formation, and revealed that macrophages were polarized towards wound healing and tumor promotion. The organoids maintained both epithelial and fibroblast lineages during the course of time, and a subset of cells in both lineages expressed the stem cell marker *Lgr5*. We identified that *Rspo3* was specifically expressed in the fibroblast lineage, providing an endogenous source of the R-spondin to activate Wnt signaling. Our studies demonstrate that air-liquid-interface-derived organoids provide a novel platform to study intercellular signaling and immune response *in vitro*.

## INTRODUCTION

Within the microenvironment of the stomach tissue, gastric epithelial cells have complex interactions with other cell lineages. These components include fibroblasts, immune lineages and endothelial cells that make up blood vessels [1]. This complex cellular milieu plays a critical role in maintaining tissue function and integrity. Similarly, solid tumor development leads to changes in the cellular microenvironment – the interactions between epithelial cancer cells and tumor stroma influence tumor development, facilitate metastasis, enable evasion of immune surveillance, and alter therapeutic responses [2]. For example, the gene expression profile from stromal cells around epithelial tumors may have prognostic implications and predictive information about response to chemotherapy [3,4]. Despite its important role in tissue regulation and maintenance, very little is known about the cellular features and intercellular communication among gastric epithelial and stromal cells.

To study the cellular microenvironment of gastric tissue, we employed an *in vitro* primary tissue culture technique that allow cells to grow and organize into three dimensional structures that resembles miniature organs, referred to as organoids. These primary tissue cultures can be derived from primary gastric tissues [5,6], epithelial stem cells [7], or induced pluripotent stem cells [8]. They are capable of self-renewal and self-organization. As demonstrated in recent studies, the organoid system enables one to maintain gastrointestinal primary tissue cultures over long periods of time as well as develop engineered cancer models where driver mutations can be systematically introduced [9-11]. For this study, organoids derived from the primary gastric tissues were grown in an air-liquid interphase **(ALI) system** [6,12]. The ALI system has a number of significant advantages compare to the conventional 2D cell culture. First, it recapitulates features of organ structure, maintains multi-lineage differentiation from the primary tissue and has the ability of self-renewal [13]. Second, one can use this method to introduce specific cancer driver events into a wild-type background, thus modeling the progression of specific oncogenes or tumor suppressors. Thus, specific cancer driver combinations can be evaluated in terms of how they contribute to oncogenesis in a stepwise manner. Third, the ALI-organoid approach enables one to study intracellular microenvironment diversity *in vitro*. This is a much closer approximation to the conditions of the original tissue compared to traditional two-dimensional cell culture. Fourth, ALI-grown gastric organoids do not require supplementation of exogenous Wnt growth factors such as Wnt3A and Rspo1 [9,11]. Rather, these primary tissue cultures are self-sustaining, suggesting that the medley of different cell lineages provides an endogenous source of growth factors.

Using a droplet-based single cell RNA sequencing **(scRNA-Seq)** technology [14-16], we analyzed thousands of individual cells and defined the gastric tissue cellular heterogeneity at single cell resolution in a self-sustaining mouse gastric organoid model. At the granularity of single cell analysis, one can derive new insights into tissue cellular heterogeneity, identify the characteristics of the diverse cell types in the local microenvironment and discover signaling interactions among different cell populations. For this study of the microenvironment, we identified different cellular lineages, determined the gene differential expression among the major cell types and as a result, identified growth factors that enable signaling crosstalk and thus maintains microenvironment cellular diversity. Based on cell type specific markers and transcriptional factors, we characterized the epithelial, fibroblast and immune cells composition of gastric organoids through the course of time and multiple passages. We found that the majority of macrophages were activated to tumor-associated M2-like status in the tissue explants microenvironment. We measured the expression of different Wnt signaling genes and determined that the growth factor, *Rspo3* demonstrated highly specific expression in mesenchymal-derived fibroblasts. Our results point to the potential role of exogenous R-spondin, as provided by the cellular microenvironment, being important for maintaining gastric epithelial cell populations.

## RESULTS AND DISCUSSION

### Single cell transcriptional profiling of primary gastric tissues and organoid cultures

Our experimental design is outlined in **Figure 1**. With scRNA-Seq, we profiled thousands of cells per sample from a series of mouse gastric tissue explants and gastric organoid cultures **(Fig 1A and Appendix Fig S1)**. For our starting material, we used a p53 null gastric tissue model originating from neonatal *Trp53*^flox/flox^ mice. Wild-type gastric organoids maintained in the ALI environment grow for only a limited time, typically 30 days and do not undergo passaging [17]. We determined that inactivation of *Trp53*, the mouse homology of *TP53*, enabled long-term culture and passages of gastric organoids in ALI environment. Moreover, TP53 loss of function is an early oncogenic transformation event, thus we replicated the start of gastric cancer development [9].

**Figure 1 -.**
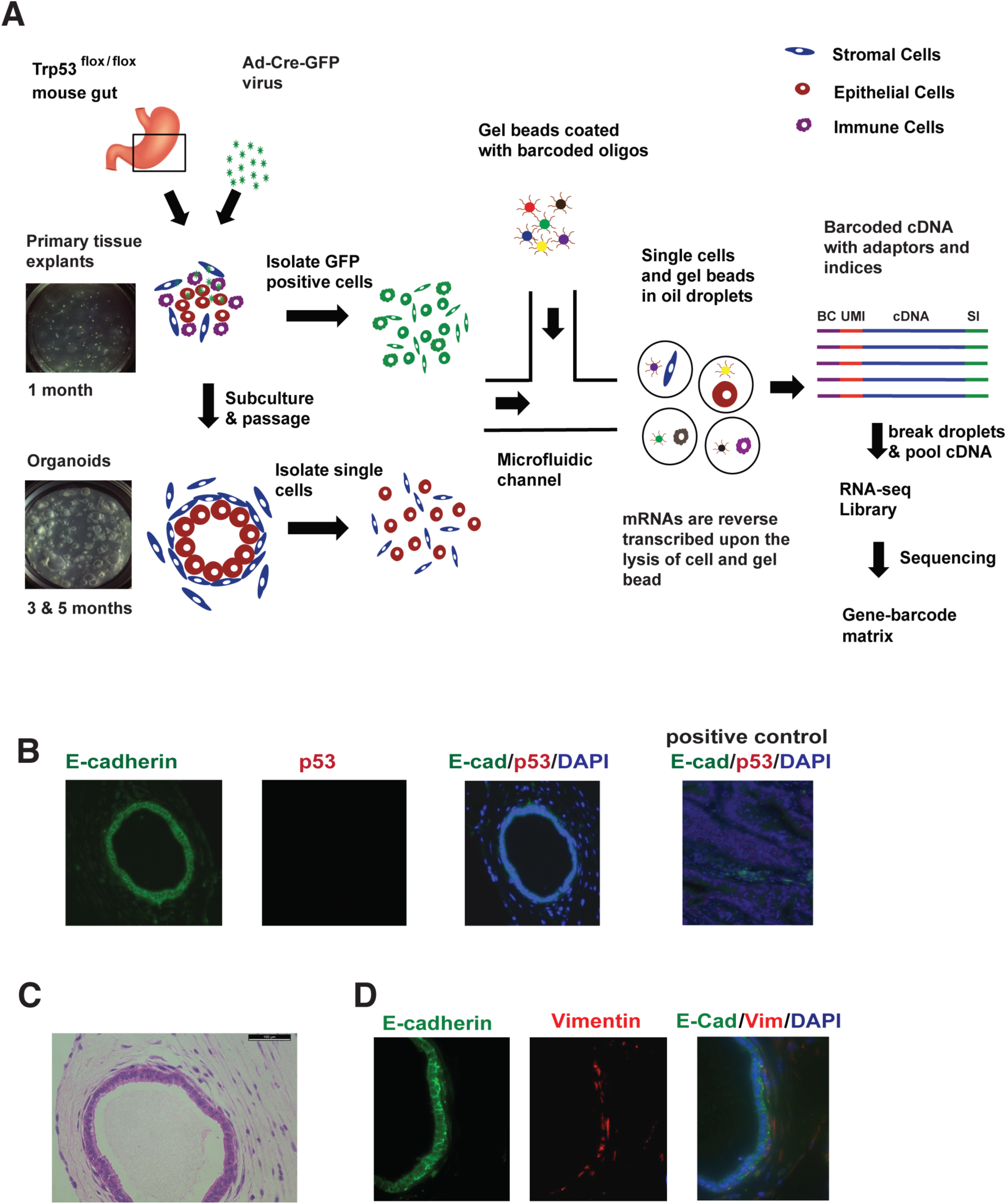
Analysis of gastric organoid populations with single cell RNA-Seq and immunofluorescence. **A** Single cells are isolated from organoids subject to scRNA-Seq **B** Loss of p53 expression in the *Trp53*-/- organoids using immunofluorescence (IF). E-cadherin is expressed in epithelial cells, and no expression of p53 was detected. A colon tumor tissue section with p53 expression is used as a positive control. Nuclei are counterstained with DAPI (blue). **C** H&E staining shows that the *Trp53*-/- organoid consist a layer of tall columnar epithelial cells with an outer lining of spindle-shaped fibroblastic stroma cells. **D** The IF showed that E-cadherin (E-cad) is expressed in epithelial cell layer and Vimentin (Vim) is expressed in surrounding fibroblast cells. Nuclei are counterstained with DAPI (blue).

Our experiments involved the following. First, we evaluated the early cell populations present in the primary gastric tissue explants after inactivation of *Trp53*. Second, we analyzed *Trp53*^*-/-*^ gastric organoids that have been cultured and passaged for three months and five months. The *Trp53*^*-/-*^ gastric organoids underwent serial passages (passage > 3) and were stably grown for more than eight weeks. We maintained these primary cultures for extended periods of time without supplements of exogenous growth factor. This result indicated the self-renewing properties of these organoid cultures.

The Cre-mediated *Trp53* deletion in gastric organoids was confirmed by genotyping **(Appendix Fig S2)**. We validated the loss of Trp53 expression with immunofluorescence **(IF)** and western blotting **(Fig 1B and Appendix Fig S3)**. Importantly, the *Trp53*^-/-^ organoid consisted an epithelial layer with surrounding fibroblastic stroma **(Fig 1C)**, which was confirmed by E-cadherin and Vimentin immunofluorescence **(Fig 1D)**.

We conducted scRNA-Seq on single cell suspensions from the gastric tissue explants and organoids. The scRNA-Seq library preparation occurred as follows: individual cells were encapsulated with single gel beads coated with oligonucleotides in droplet partitions via a high throughput microfluidic device [15]. Each oligonucleotide is consisted of a 30nt poly-A primer, a 14nt cell barcode, and a 10nt random sequence as unique molecular identifier **(UMI)** to eliminate molecular duplicates and enable single molecule transcript counting. Upon the lysis of cell and gel bead, mRNAs were reverse transcribed by the barcoded oligonucleotides in individual droplets. Subsequently, the droplets were broken and barcoded cDNAs were pooled together for PCR amplification to generate complete scRNA-Seq libraries for sequencing.

In total, we sequenced 5,822 cells from three samples: (1) 2,304 cells from the *Trp53*^*-/-*^ gastric tissue explants in culture for one month; (2) 2,087 cells from the *Trp53*^*-/-*^ organoids in culture for three months (passage = 4); (3) 1,710 cells from the *Trp53*^*-/-*^ organoids in culture for five months (passage = 6) **(Table 1)**. To guarantee mRNA transcripts were adequately sequenced for distinguishing the cell types, we generated more than 200 million reads for each sample, and more than 90,000 reads per cell. A previous study has shown that 50,000 reads per cell is sufficient for accurate cell-type classification and biomarker identification [18]. The median number of genes and mRNA transcripts (UMI counts) detected per cell were higher in the gastric tissue explants cells (∼2900 and ∼11000) compared to the cells from two organoid samples (∼2000 and ∼5000) **(Table 1 and Appendix Fig S4)**.

**Table 1.**
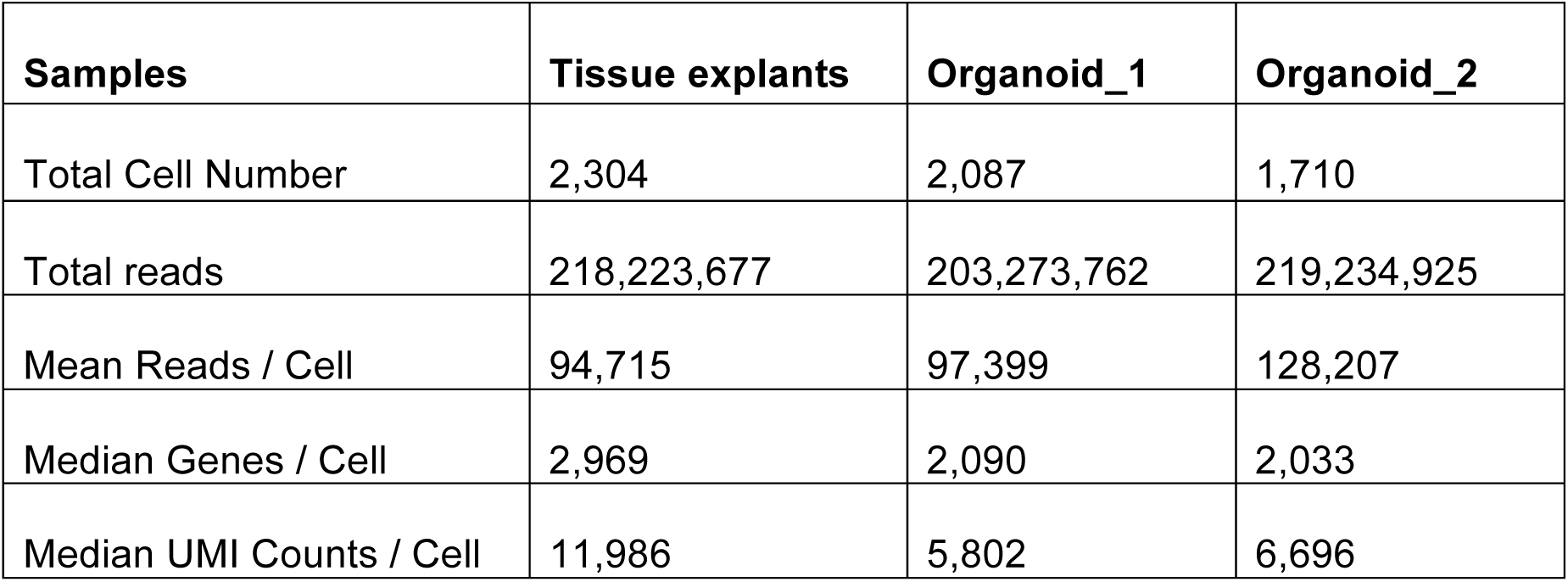
Single cell sequencing metrics for gastric organoids.

We used the program Seurat to analyze the data [19]. As a quality control metric, we evaluated genes that were expressed in three cells or more and included those cells that had 500 gene transcripts or more. To remove potential cell doublets where two cells exist within a single droplet partition as well as low quality cells with poor RNA-Seq data [20], we filtered out those cells that have unique gene counts over 5,000 or contain more than 5% the percentage of mitochondrial genes **(Appendix Fig S5)**. Principal component analysis **(PCA)** was performed on remaining cells to reduce the dimensionality of the scRNA-Seq data matrix using high variable genes [16]. Subsequently, cells were clustered based on a graph-based clustering approach [21,22] and were visualized in two dimensional space using t-distributed Stochastic Neighbor Embedding **(tSNE)** [23].

### Characterization of single cells from primary gastric tissue

Our analysis indicated that gastric tissue explants were undergoing macrophage-based tissue remodeling in ALI. Five cellular clusters were apparent on the tSNE map **(Fig 2A)**. To identify cluster specific genes, each cluster of cells was compared to all other cell clusters by differentially expression analysis **(Table EV1)**. Clustering of specific genes revealed three main cell populations in the primary tissues **(Fig 2A)**: fibroblast cells (cluster 1), epithelial cells (cluster 2), and leukocytes (clusters 3-5). Fibroblast specific genes, such as *Col1A1*, *Bgn*, *Dcn* [24], were significantly expressed in the cluster 1 **(Fig 2B)**, while epithelial specific markers, such as *Epcam*, *Krt7* and *Krt14* [25], were significantly expressed in cluster 2 **(Fig 2C)**. Most of the cells in clusters 3-5 expressed leukocyte common antigen *Cd45* (*Ptprc*) **(Fig 2D)**. Moreover, the expression of *Cd115* (*Csf1r*), *Cd11b (Itgam), and Cd18* (*Itgb2*) in clusters 3-5 indicated that these cells were monocytes derived tissue macrophages **(Fig 2E)** [26].

**Figure 2 -.**
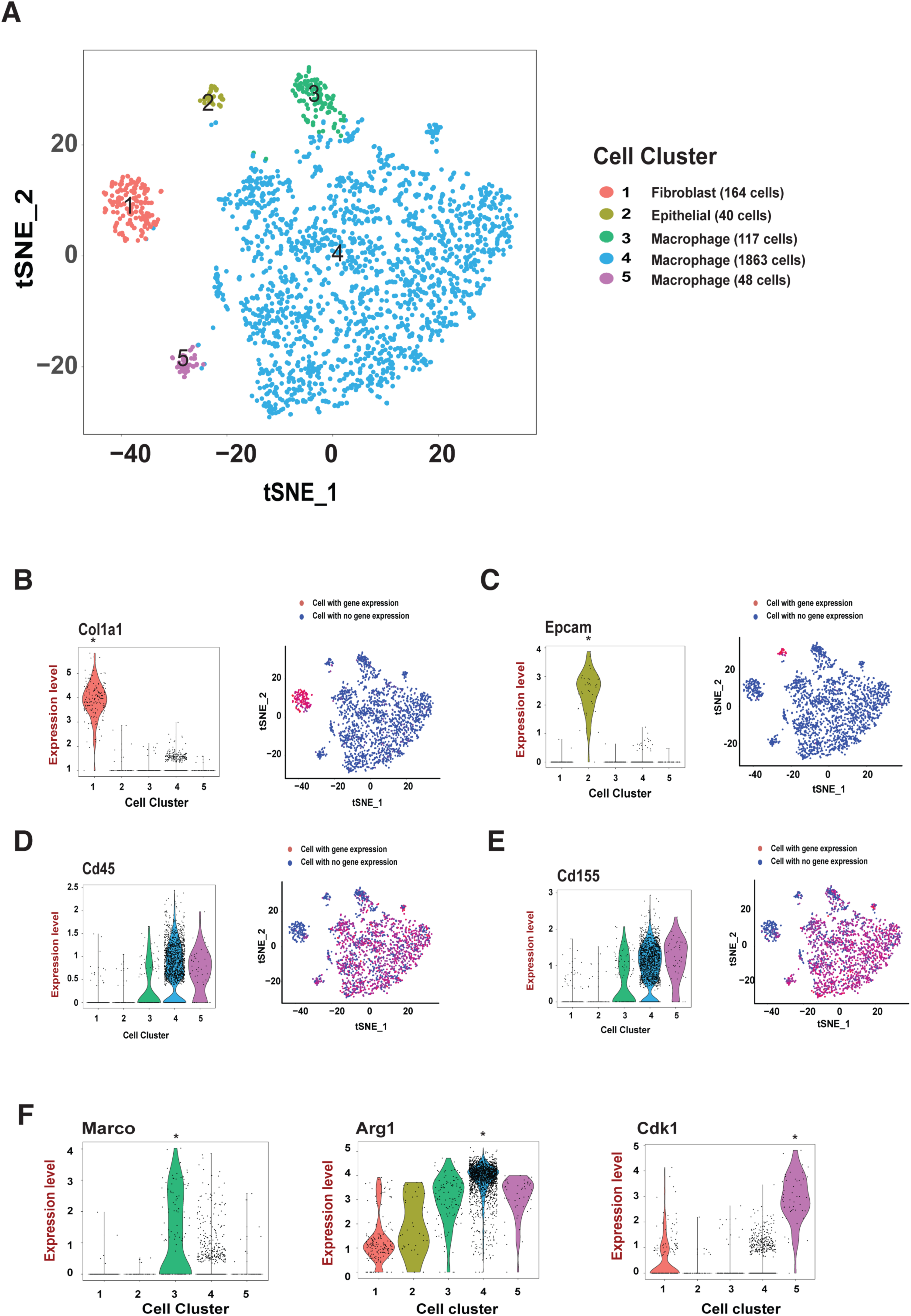
Distinct cell types in the tissue explants. **A** tSNE projection of 2,232 cells from tissue explants. Cells are grouped into five clusters based on transcriptome profiles and are colored accordingly. The cell type assignment of each cluster is based on the gene expression analysis. **B-E** The expression of cluster specific genes, i.e., *Col1a1*, *Epcam*, *Cd45*, *Cd155*, are displayed on violin plots and tSNE maps. **F** The violin plots of macrophage group specific genes, i.e., *Marco*, *Arg1*, *Cdk1*. Data information: In (**B-F**), every dot represents an individual cell in both violin plots and tSNE maps. The gene expression level of the natural log of the normalized UMI counts in the violin plot. * Bonferronii adjusted P value <0.001.

To evaluate the three macrophage clusters, we performed gene ontology **(GO)** enrichment analysis using the EnrichR program [27], and identified the top ranked genes. There were distinct gene expression patterns among the macrophages: cluster 3 was enriched with the inflammatory responses and cell clearance related genes, cluster 4 was enriched with the leukocyte cytokine and migration genes, and cluster 5 was enriched with cell cycle genes **(Table 2)**. Our results suggested that a subgroup (∼6%) of the macrophages (cluster 3), expressing scavenger receptor Marco and cytokine Il18 [28], and thus, had inflammatory characteristics consistent with what is described as the M1 class **(Fig 2F)**. Cluster 4 demonstrated an upregulation of genes associated such as *Arg1* (anti-inflammatory) and *Mmp12* (extracellular matrix remodeling) [29]. These genes are expressed in tumor-activated macrophages **(TAMs),** also known as M2 macrophages, a category of immune cells that play a critical role in tissue remodeling. Cell cycle genes, such as *Cdk1, Aurkb* and *Ccna2* were significantly upregulated in cluster 5, indicating a subgroup (∼2%) of macrophages was actively proliferating. This finding is in agreement with previous studies where it was observed that tissue macrophages undergo cell division within the primary tissue where they reside [30]. At this early explant stage, our ability to identify macrophage types and the increased number was related to isolating *Trp53*^*-/-*^ single cells using GFP signals from tissue explants. The cells were infected by the Ad-Cre-GFP therefore the GFP signals were more likely to remain in the non-proliferating, long half-life tissue-resident macrophages than other cell lineages.

**Table 2.** The top five GO biological processes in each cluster based on the gene enrichment analysis.

|  |  |
| --- | --- |
| **M1-like Macrophage (Cluster 3)** | **Adjusted p-value** |
| Regulation of superoxide anion generation (GO:0032928) | 0.000819 |
| Endocytosis (GO:0006897) | 0.000819 |
| Apoptotic cell clearance (GO:0043277) | 0.001645 |
| Regulation of superoxide metabolic process (GO:0090322) | 0.001891 |
| Regulation of response to wounding (GO:1903034) | 0.004406 |
| **M2-like Macrophage (Cluster 4)** | **Adjusted p-value** |
| Leukocyte chemotaxis (GO:0030595) | 2.43E-09 |
| Leukocyte migration (GO:0050900) | 5.45E-08 |
| Cell chemotaxis (GO:0060326) | 5.45E-08 |
| Chemotaxis (GO:0006935) | 1.43E-07 |
| Taxis (GO:0042330) | 1.43E-07 |
| **Proliferating Macrophage (Cluster 5)** | **Adjusted p-value** |
| Mitotic cell cycle (GO:0000278) | 2.32E-28 |
| Nuclear division (GO:0000280) | 4.70E-15 |
| Organelle fission (GO:0048285) | 1.24E-14 |
| Mitotic nuclear division (GO:0007067) | 1.07E-13 |
| Chromosome organization (GO:0051276) | 2.45E-12 |

Top 50 genes in each cluster were included in the enrichment analysis. P value was calculated with the Fisher's Exact Test and adjusted by the Bonferroni correction [27].

In conclusion, we found that the majority of macrophages were activated to tumor-associated M2-like status and only a small population was similar to inflammatory M1-like. This result suggests that the macrophages in the initial tissue explant microenvironment were polarized towards wound healing and tumor promotion. Importantly, this result indicates that the ALI organoid method may enable one to manipulate and study the microenvironment consequence of tumor-associated macrophages *in vitro*.

### Gastric organoid microenvironment has differentiated cellular lineages

We identified multiple cellular lineages from stable gastric organoid cultures maintained over time. In particular, we observed gene expression patterns at single cell resolution that had the properties of primary gastric tissue. For this analysis, we sequenced 2,087 cells from the three-month old culture that we refer to as Organoid_1 (culture time = three months, passage = 4). A total of 1,961 cells had adequate sequencing quality and were grouped into two clusters on the tSNE map **(Fig 3A)**. Two major lineages, epithelial and mesenchymal-derived fibroblasts, were clearly denoted based on gene expression patterns and clustering **(Fig 3B-D and Table EV2)**. The epithelial cell type was enriched with markers, such as *Epcam*, *Krt7* and *Krt14* and the fibroblast type was enriched with lineage-specific markers, such as *Col1a1*, *Bgn* and *Dcn*. Unlike the gastric tissue explants, cells from the primary organoid cultures did not express leukocyte lineage markers or macrophage markers – this result indicated that the period of tissue remodeling had been completed and hematopoietic cells could not be continuously subcultured despite initial presence in the ALI environment. A notable finding is the lack of expression of intestinal lineage markers, such as *Muc2*, *Cdx1* and *Cdx2*, indicating that these cells were strictly of gastric origin.

**Figure 3 -.**
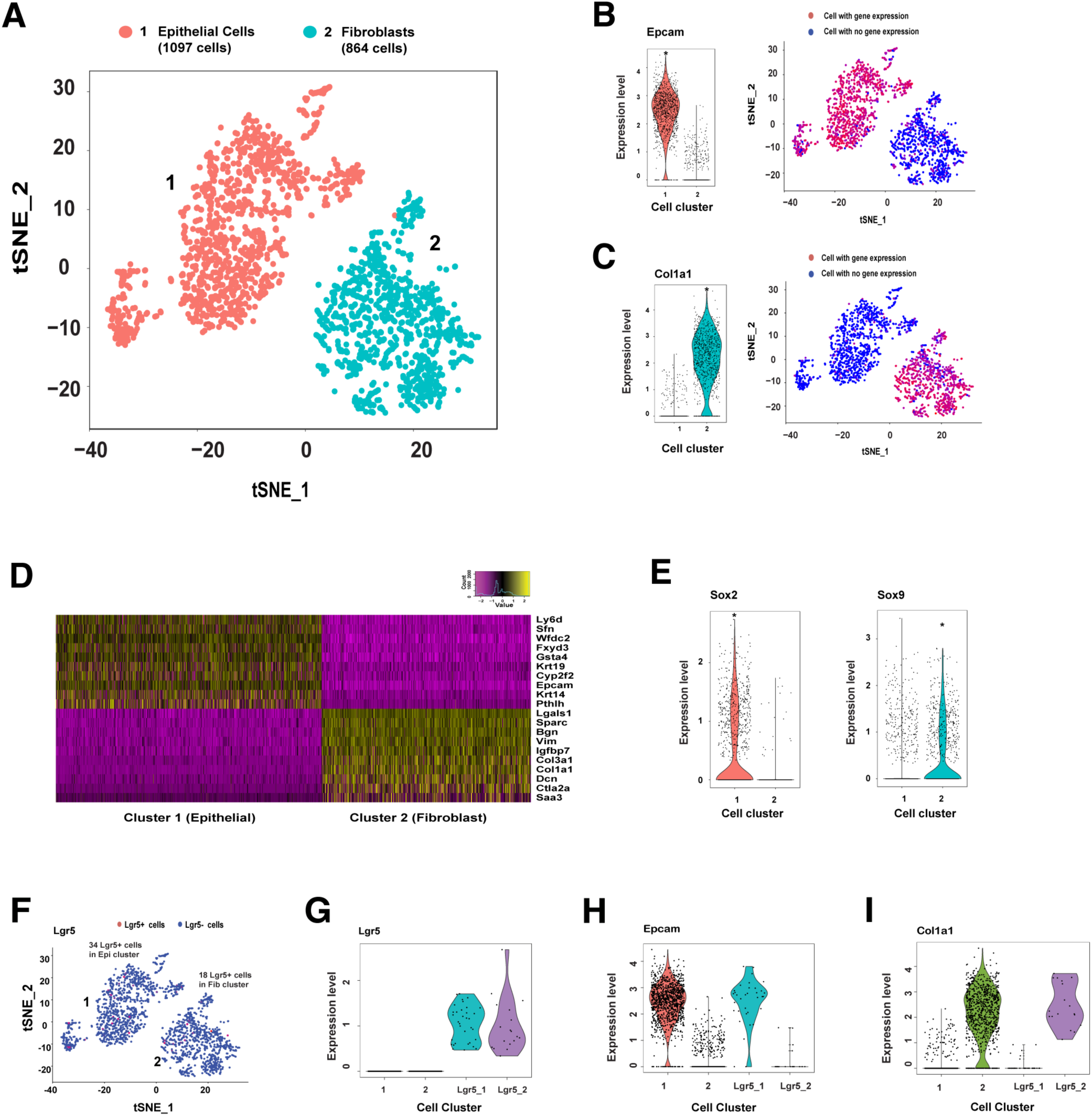
Organoids are composed of epithelial and fibroblast cell types. **A** tSNE projection of 1961 cells from Organoid_1. Cells are grouped into two clusters, epithelial and fibroblast, based on transcriptome profiles and were colored accordingly. **B-C** The expression of cluster specific genes, *Col1a1* and *Epcam,* are displayed on violin plots and tSNE projection. **D** The heatmap of top ten genes associated with each cluster. Each row represents a gene, and each column represents a single cell. **E** Transcriptional factors, *Sox2* and *Sox9*, are expressed in the epithelial and fibroblast clusters respectively. **F** tSNE plot shows that *Lgr5* mRNAs present in both epithelial and fibroblast cell clusters. Epi – Epithelial, Fib – Fibroblast. **G** *Lgr5* expression in Lgr5+ only epithelial (Lgr5_1) and fibroblast (Lgr5_2) cell clusters. The remaining epithelial and fibroblast cells form the cluster 1 and 2 respectively. **H-I** The expression of epithelial and fibroblast lineage markers, *Col1a1* and *Epcam,* in Lgr5+ cells. Data information: The expression level of violin plot is the natural log of the normalized UMI counts. * Bonferronii adjusted P value <0.001

We evaluated the expression of key transcriptional factors and determined the development status of these tissues **(Fig 3E and Appendix Fig S6)**. The epithelial cells expressed transcriptional factors *Sox2* and *Klf5*, which are typically expressed in progenitor cells in the immature epithelium of the gut [8]. The majority of cells in the epithelial cluster expressed forkhead transcription factor *Foxq1*, which is specifically expressed in mucin-producing foveolar cells of stomach lineage [31]. We also detected a small number of cells that express ghrelin (*Ghrl*), indicating the presence of endocrine cells. Similarly, the fibroblast cells expressed *Sox9* and *Hoxa5*, two transcriptional factors that are specifically expressed the undifferentiated cells in the mesenchyme of immature gut [32]. Overall, these results suggest that these gastric organoids had cellular properties of the mouse stomach.

### Stem cell lineages present in organoids

Stem cells provide self-renewal of organoids. Studies have shown that single Lgr5+ cells were capable of replenishing gastric cells, and thus is a marker of gastric stem cells [13]. We found that ∼3% of cells (34 cells) in the epithelial cluster express *Lgr5* mRNA **(Fig 3F)**. Interestingly, ∼2% of cells (18 cells) in the fibroblast cluster also expresses *Lgr5* mRNA **(Fig 5F)**. A previous study only observed Lgr5 mRNA expression among wild-type gastric epithelial cells, but not in the fibroblast cells [33]. However, a recent study found LGR5 was expressed in both epithelium and stromal cells in normal colon tissue, and the expression level was notably increased in tumor samples [34]. To further characterize the heterogeneity between Lgr5+ cell lineages, we selected and generated the Lgr5+ only cell clusters with epithelial (Lgr5_1) and fibroblast (Lgr5_2) lineages from the respective clusters **(Fig 3G)**. Per single cell gene expression analysis, epithelial- and fibroblast-specific genes distinguished the two Lgr5+ groups. For example, *Epcam* and *Dcn* were significantly upregulated in Lgr_1 cells, while *Col1a1* and *Krt14* were significantly upregulated in Lgr5_2 cells **(Fig 3H-I and Table EV3)**. Our study suggests that both epithelial and fibroblast lineage cells express Lgr5 mRNA within a stem cell population.

### Gastric organoid transcriptional phenotype is stable over time

We sequenced and analyzed 1,710 cells from a second organoid sample (Organoid_2) that was cultured for five months (passage = 6). Indicating the reproducibility of our findings, the single cell gene expression profiles of both Organoid_1 and Organoid_2 were very similar as noted by a linear regression comparison for the fold change expression levels (r = 0.83, P <0.001) **(Table 1 and Appendix Fig S7)**. Organoid_2 cells were grouped into two distinct epithelial and fibroblast cell clusters after PCA and tSNE analysis **(Appendix Fig S8)**. Subsequently, our cluster analysis revealed that the transcriptional profiles from Organoid_2 cells highly correlated with Organoid_1 cells. Namely, our analysis of both data sets identified the same lineage types **(Appendix Fig S10 and Table EV4)**. Again, we observed the same cell type specific expression of transcriptional factors, such as *Sox2*, *Sox9*, and *Foxq1*. Furthermore, 43 cells (2.5%), from both epithelial and fibroblast lineages, expressed *Lgr5* mRNA in the Organoid_2, thus providing another set of independent results point to the presence of a stable stem cell population. Similar to the three-month culture (Organoid_1), the organoid cells did not express hematopoietic lineage markers. Overall, these results indicated that the cellular diversity of the gastric organoids was stably retained through time.

### Distinct Wnt/beta-catenin signaling activation between the cellular lineages

Wnt/b-catenin signaling pathways regulates the homeostasis of gastrointestinal tract and maintain the self-renewal capability of epithelial stem cells [35]. Many methods for primary tissue cultures only maintain pure epithelial cells. To sustain their growth in tissue culture, these isolated epithelial cells require external supplementation of niche factors such as Rspo1, Wnt3A, Noggin [5] or external fibroblasts as feeder cells [36]. Given that ALI-maintained, multi-lineage organoid cultures did not require external supplementation, we hypothesized that secretion of endogenous niche factors by fibroblast cells in stroma sustained Wnt/β-catenin signaling and stem-cell renewal in organoids.

To examine the effect of Wnt inhibition, we treated the gastric organoids with the Wnt signaling inhibitor LGK974. This molecule targets Porcupine, a Wnt-specific acyltransferase and thus, inhibits Wnt signaling *in vitro* and *in vivo*. Previous studies have shown that, inhibition of Wnt signaling leads to cessation of epithelial cell proliferation and loss of crypt in mouse as well as in organoids derived from primary intestinal tissues [6-37,38]. After LGK974 treatment, the organoid epithelial layer degenerated and there was notable growth inhibition as observed with Ki67 staining **(Appendix Fig S9)**. These results indicated that this gastric organoid model, despite p53 inactivation, recapitulated the *Wnt* niche dependency *in vitro*.

We examined our results from scRNA-Seq to define the Wnt dependencies among the major cell types. Among 394 Wnt related genes as defined by gene ontology, 330 were expressed among the Organoid_1 cells **(Table EV5)**. Thirty genes were differentially expressed with statistical significance **(corrected p< 0.05 and fold change > 1.5)** between the epithelial and stromal epithelial cell populations **(Fig 4A)**. Notably, these genes included specific cell lineage markers implicated in the Wnt signaling pathway, such as *Col1a1*, *Krt6a* and *Cdh1*. Highly expressed in fibroblast cells were the stomach mesenchymal transcription factor *Barx1* and its downstream targets *Sfrp1* and *Sfrp2*, two genes that function as Wnt antagonists. Studies have shown in gastric mesenchymal cells, Barx1 signals through Sfrp1 and Sfrp2 to control stomach-specific epithelial differentiation [39].

**Figure 4 -.**
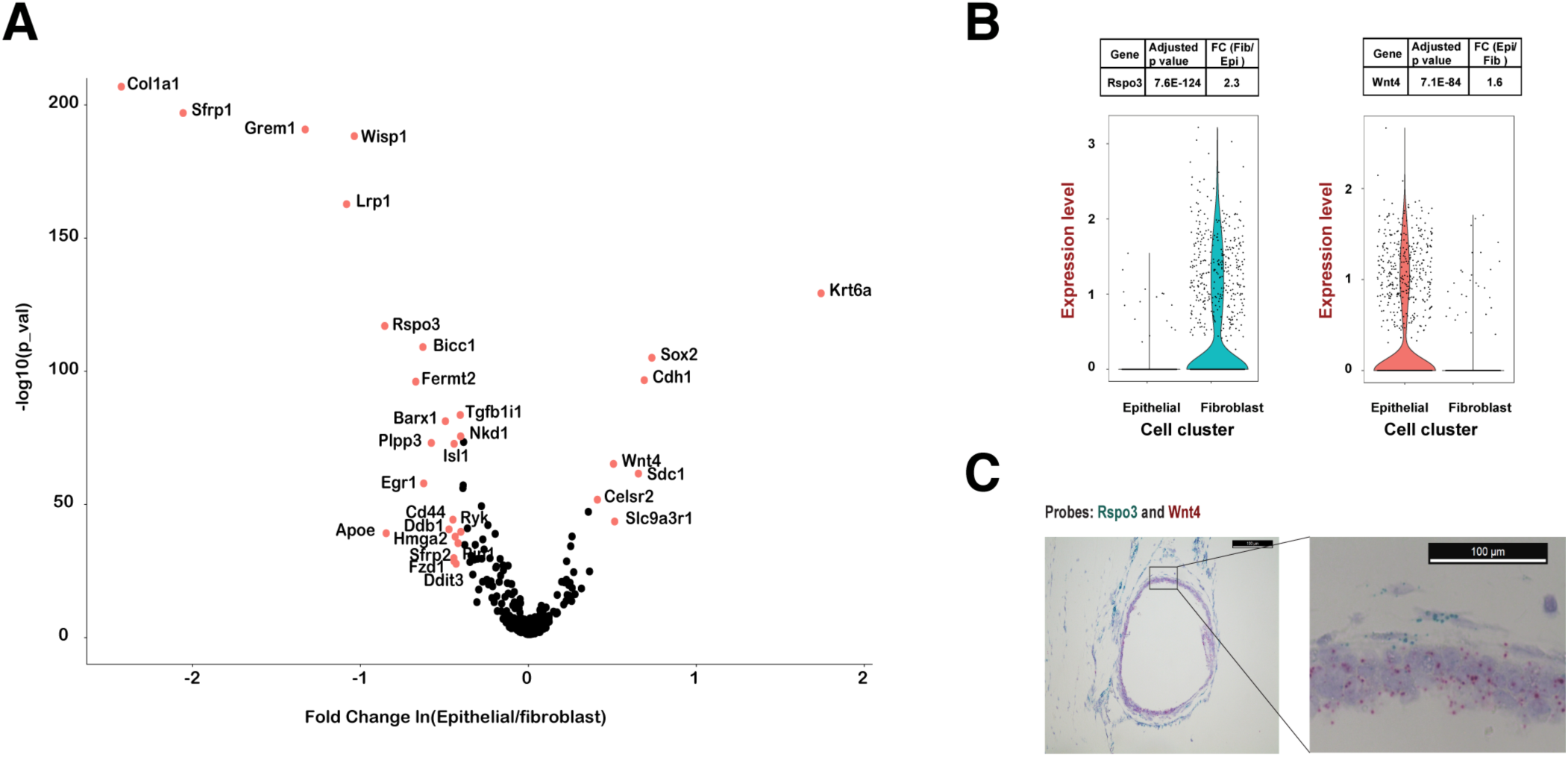
Distinct Wnt signaling activation and ligand expression between epithelial and fibroblast cells. **A** Differentially expressed Wnt related genes between epithelial and fibroblast cells. Genes with fold change > 1.5 (|Ln| >0.4) and Bonferonii adjusted P< 0.05 were labeled in red with gene name. The x-axis is the natural log transformed fold change of gene expression in epithelial vs mesenchymal cells. The y-axis is the −log10 transformed p value. **B** Cell type specific expression of *Rspo3* and *Wnt4*. The violin plots depict that *Rspo3* mRNA expression is enriched in the fibroblast cells and *Wnt4* mRNA expression is enriched in the epithelial cells. The expression level of violin plot is the natural log of the normalized UMI counts. **C** RNA In situ hybridization confirms cell type specific expression of *Rspo3* and *Wnt4* in the organoid. The organoid FFPE section is hybridized and amplified by the probes specifically targeting *Rspo3* and *Wnt4* mRNAs. Rspo3 probes are the blue dots detected in the fibroblast cells surrounding the organoid, and Wnt4 probes are the red dots detected in the epithelial cells forming the wall of the organoid sphere (Scale bar = 100uM).

### Lineage-specific expression of *Rspo3* and *Wnt4* suggest niche crosstalk

Wnt proteins bind to a variety of different receptors that include FZD, LRP5 and LRP6. This interaction leads to β-catenin nuclear translocation and Wnt/b-catenin activation. Similarly, the family of R-spondin growth factors (RSPO1-4) interact with LGR4-LGR6 receptors, inhibit the degradation of FZD and potently activate Wnt/b-catenin signaling [40]. However, Wnt and R-spondin proteins are not equivalent in terms of their function [36]. Unlike R-spondin ligands, Wnt proteins cannot induce Lgr5+ stem cell self-renewal, but instead prime the expression of R-spondin receptors. Given the differential expression of Wnt genes among the various lineages, we examined gene expression for secretory factors between the two major cell populations.

Based upon single cell analysis, we found that the gene expression of *Rspo3* was significantly enriched in the fibroblast cells, while the gene expression of *Wnt4* was significantly enriched in the epithelial cells **(Fig 4B)**. The lineage-specific expression of *Rspo3* and *Wnt4* was consistent in both long-term gastric organoid cultures **(Appendix Fig S10)**. Thus, this result was reproducible. For the initial gastric tissue explants, we observed specific expression of *Rspo3* in the fibroblast cell. However, the expression of *Wnt4* was very low in the epithelial cells (**Appendix Fig S10**).

We confirmed that the *Rspo3* gene was specifically expressed in the fibroblast cells and *Wnt4* was specifically expressed in the epithelial cells. As an orthogonal validation, we applied RNA *in situ* hybridization **(ISH)** to directly examine mRNA expression in organoid cells. First, we evaluated the sensitivity and specificity of the RNA ISH with two positive controls. Specifically, we measured the expression of two housekeeping genes *Ppib* and *Polr2a* and found that they were simultaneously expressed in all organoids cells while there was no expression of a negative control, the E.coli gene *dapB* gene **(Appendix Fig S11)**. Using gene-specific probes, we confirmed that *Wnt4* mRNA was specifically expressed in the epithelial cellular layers making up the cystic portion of the organoid. In contrast, the *Rspo3* mRNA was specifically expressed the adjacent lining composed of fibroblast cells **(Fig 4C)**. Overall, we confirmed the lineage specificity of expression based on ISH localization of the *Wnt4* and *Rspo3* transcripts. Our results implicated *Rspo3* and *Wnt4* are candidates for mediating intercellular communication and maintain different cell types in gastric tissues. Importantly, we identified Rspo3 as the endogenous source of Rspos supplied by fibroblasts in the gastric stroma. Recently, Virshup et al. found that Rspo3 to be the predominant Rspos expressed in cultured mouse intestinal stroma and was sufficient to support the intestinal homeostasis [36]. More recently, another report noted that stromal Rspos sustain gastric epithelial stem cells and gland homeostasis [41]. Together, our studies suggested that fibroblasts are the likely source of Rspos that support the growth niche in gastrointestinal TME. Recurrent RSPO fusions were identified in a subset of colon cancers and were mutually exclusive with APC mutations, suggesting that they are involved in the activation of Wnt signaling [42]. The RSPO fusions probably render epithelial cells to lose the dependency on the RSPO3 secreted by the fibroblasts. More recently, targeting RSPO3 in colon tumors with RSPO fusions promoted differentiation and loss of stem-cell function [43]. Our data suggests that therapeutic strategies targeting RSPO3 may be relevant to gastric cancers of epithelial origin.

Taking into account the potential transcriptional consequences of *Trp53*^*-/-*^ model, we conducted an scRNA-Seq study on neonatal gastric tissue explants with wild-type *Trp53*. Overall, we observed the similar lineage-specific expression of *Rspo3* and *Wnt4*, suggesting the expression of Wnt ligands were independent of p53 status. For this comparison we sequenced 955 cells from Trp53 wild-type gastric explant at average 189,775 reads per cells, and 869 cells passed the QC. We observed two major cell lineages, epithelial and mesenchymal fibroblast cell types formed distinct clusters, and were enriched with the same lineage specific markers such as *Epcam*, *Krt7*, *Col1a1*, and *Dcn*, as noted in the p53 null organoids. Among Wnt related genes, we found similar lineage specific activation, such as *Krt6a* (adjust p value = 4.0 E-41) in epithelial cells and *Col1a1* (adjust p value = 2.5 E-16) in fibroblasts. Again, we found *Rspo3* was significantly enriched in the fibroblast cells (adjusted p value = 1.3 E-43), and the expression of Wnt4 was higher in epithelial cells despite very low expression in both cell types, mirrored the expression pattern in the p53 null gastric explant. These results support our conclusion that p53 loss of function does not alter the lineage specific transcriptional profiles.

In summary, our studies systematically characterized primary gastric tissues using organoids composed of epithelial cells and stromal cells in single cell-resolution. We demonstrated that this system could recapitulate the gastric tissue microenvironment and intercellular signaling *in vitro*. Incorporating this methodology in cancer biology will allow us to culture tumor with its stromal components, providing a novel tool to examine the signaling crosstalk between tumor cells and associated stromal cells. Moreover, it can further extend into a screening platform for drugs targeting tumor microenvironment.

## MATERIALS AND METHODS

### Organoid development and growth

The Stanford University Administrative Panel on Laboratory Animal Care approved all animal experimental protocols. *Trp53*^flox/flox^ mice were kindly provided by Dr. Anton Berns (Meuwissen, Linn et al. 2003). We dissected stomachs from neonatal mice (age P4-7) and washed them in cold F12 to remove all luminal contents. The neonatal stomach was extensively minced and embedded in collagen gel using a double-dish culture system as previously described [6]. Cre recombinase adenoviruses (Ad-Cre-GFP, Vector Biolabs) were added the cultures to induce *Trp53* deletions in the cultured gastric tissue. The cultures were checked for GFP signaling by fluorescence microscopy after 3 days to confirm the infection. We replace organoid growth media (F-12 nutrient mixture, 20%FBS, 1%Antibiotic-Antimycotic) every week. The organoids were passaged at a 1:2 ratio every three to four weeks. To determine the effects of Wnt inhibition on these organoids, we treated the organoid culture with 10 uM LGK974 (Selleck Chemicals) for a period of 7 days.

### Molecular characterization of mouse gastric organoids

The mouse genomic DNA (gDNA) was extracted from the tail biopsy using the Maxwell® 16 Tissue DNA Purification Kit (Promega), and the organoids gDNA was extracted using QuickExtract™ DNA Extraction Solution (Epicentre). The amount of DNA was quantified by Qubit® dsDNA BR Assay Kit (Invitrogen). The PCR primer sequences used for genotyping: Trp53_Fw: CACAAAAACAGGTTAAACCCAG. Trp53_Rv: AGCACATAGGAGGCAGAGAC. Cdh1_Fw: GGGTCTCACCGTAGTCCTCA. Cdh1_Rv: GATCTTTGGGAGAGCAGTCG. The PCR was performed using the Q5® Hot Start High-Fidelity 2X Master Mix (NEB) according to the manufacturer’s protocol. Briefly, initial denaturation at 98 °C for 30 sec, followed by 30 cycles of amplification (98 °C for 10 sec, 64 °C for 10 sec, 72 °C for 15 sec), and a final extension at 72 °C for 3 min. The PCR products were run on 2% agarose gel.

### Western blotting

Cell lysates (30 µg) were separated on 4–15% precast polyacrylamide gels (Mini-PROTEAN® TGX™ Precast Protein Gels, Bio-Rad) and were transferred to nitrocellulose membranes. The antibodies used for blotting included p53 (Santa Cruz, sc-1311-R) and beta actin (Abcam, ab8227).

### Immunofluorescence and immunohistochemistry studies

The organoid samples were fixed with 4% paraformaldehyde overnight and paraffin-embedded as previously described [6]. Sections (∼5 uM) from blocks were deparaffinized and stained with H&E for histology analysis. For IF staining, we used primary antibodies for the following proteins: p53 (1:100, Santa Cruz, sc-1311-R), E-cadherin (1:300, BD Biosciences, 610182) and Vimentin (1:100, Cell signaling, 5741). The secondary antibodies used were Alexa Fluor 488 goat anti-mouse (1:500; Invitrogen, A10680) and Alexa Fluor 555 goat anti-rabbit (1:500; Invitrogen, A21428). The ProLong Gold Antifade Reagent with DAPI (Cell Signaling) was used for mounting. For IHC staining, we used Ki67 (1:200, Cell signaling, 9027), SignalStain® Boost IHC Detection Reagent (Cell signaling, HRP, Rabbit) and SignalStain® DAB Substrate Kit (Cell signaling).

### Organoid disaggregation, single cell library preparation and flow cytometry

The gastric organoids were isolated from collagen gel by incubating with collagenase IV (500mg/mL, Worthington) in a 15-mL Falcon tube at 37 °C for up to 1 hour. The tube was centrifuged at 400g for 5 min. The supernatant was discarded and the organoid pellet was washed twice with 10 mL F-12. The organoids were collected by centrifuging and were re-suspended with 500 uL Trypsin-EDTA (0.25%, Gibco) at 37 °C for 25 min to dissociate into single cells. We used a P200 Pipette to pipet the cell suspension up and down several times to ensure digestion. The gastric cells were washed with 5 mL organoid growth media (F-12 nutrient mixture, 20% FBS), and filtered twice through cell strainers (mesh size: 40 uM). After being centrifuged at 400 g for 5 min, the supernatant was discarded, cells were washed with 1X PBS and were re-suspended at ∼1000 cells/uL in 1X PBS containing 0.4% BSA. The GFP signal was detectable in tissue explants 1 month after the Ad-Cre-GFP infection and individual cells with positive GFP signal were isolated using a BD Influx cell sorter.

The single cell RNA-Seq libraries were prepared using the 10X Genomics Single Cell 3’ Gel Bead and Library Kit following the manufacturer’s instruction. Briefly, cell suspensions were loaded on 10X Genomics Single Cell Instrument where single cells are partitioned in droplets. Upon encapsulation, cells are lysed and the gel bead dissolution releases sequencing adapter oligonucleotides that mediated the reverse transcription of poly-adenylated RNAs. With the incorporation of the oligonucleotide adapter, each cDNA molecule incorporates a UMI and droplet partition barcode. Subsequently, the emulsions were broken and barcoded cDNAs were pooled for PCR amplification. Amplified cDNAs were shared to ∼ 200bp using a Covaris E210 system. After end repair and A-tailing, adapters were ligated to the sheared DNA product, followed by PCR to incorporate sample indices. The scRNA-Seq libraries were run on an Agilent High Sensitivity DNA chip for quality control, and were quantified using the KAPA Library Quantification Kits for Illumina® platform (KAPA Biosystems). The libraries were loaded at 20 pM on an Illumina HiSeq 2500 using the TruSeq v3 200 cycles HS kit or at 1.8 pM on an Illumina NextSeq 500 using the 150 cycles High Output kit. Standard Illumina paired-end sequencing with dual indexing uses the following read length: Read 1 - 98bp (RNA read), Read 2 - 10bp (UMI, tissue explant library only have 5bp), i7 Index – 14 bp (Single cell index), i5 index – 8bp (Sample barcode).

### Data analysis

The Chromium Single Cell Software Suite (http://support.10xgenomics.com/single-cell/software/overview/welcome) was used to demultiplex samples, process barcodes, and count single cell genes [44]. Briefly, FASTQ files were generated from Read 1 (RNA read), Read 2 (UMI) and i7 index (single cell index) after sample demultiplexing based on i5 index. Read 1 was aligned to mouse reference genome mm10 using STAR [45]. Subsequently, single cell index and UMI were filtered using the Cell Ranger pipeline [44]. The final output was a gene-barcode matrix contains only confidently mapped (MAPQ = 255), non-PCR duplicates with valid barcodes and UMI. Matrices from multiple libraries were combined together by concatenation in the Cell Ranger R Kit (<u>http://support.10xgenomics.com/single-cell/software/pipelines/latest/rkit</u>).

The gene-barcode matrices were further analyzed and visualized based on the Seurat R package [19]. For quality control, the gene expressed less than 3 cells and the cell contained less than 500 genes were discarded. To remove potential cell doublets and low quality cells [20], the cells that have unique gene counts over 5,000 or the percentage of mitochondrial genes more than 5% were filtered out.

Top variable genes across single cells were identified using the method described in Macosko et al. [16]. Briefly, the average expression and dispersion were calculated for each gene, genes were subsequently placed into 20 bins based on expression, and then calculating a z-score for dispersion within each bin. Principal component analysis (PCA) was performed to reduce the dimensionality on the log transformed gene-barcode matrices of top variable genes [16]. Cells were clustered based on a graph-based clustering approach [21,22], and were visualized in 2- dimension using tSNE [23]. Likelihood ratio test that simultaneously test for changes in mean expression and in the percentage of expressed cells was used to identify significantly differentially expressed genes between clusters [46].

### In situ RNA hybridization

RNA in situ hybridization was performed on organoid FFPE tissue sections using the RNAscope® 2-plex Reagent Kit (Advanced Cell diagnostics) following the manufacturer’s instructions. Briefly, the FFPE tissue sections were deparaffinized, boiled in the target retrieval reagent for 8 mins and digested by protease for 15 mins, followed by a series of probe hybridization and signal amplification. The slides were counterstained in 50% hematoxylin for 30sec, mounted with VectaMount (Vector laboratories), and examined under a bright field microscope. The RNAscope probes used were as follows: Mm-Rspo3 (402011), Mm-Wnt4-C2 (401101-C2), RNAscope® 2-plex positive control probe_Mm (320761), and RNAscope® 2-plex negative control probe (320751).

### Data deposition

The sequencing data from this study were deposited at NCBI Sequence Read Archive (Accession number: SRP104455).

## Acknowledgements

J.C. and H.P.J. were supported by a Research Scholar Grant, RSG-13-297-01-TBG from the American Cancer Society. H.P.J. received additional support from the Doris Duke Clinical Foundation Charitable Foundation Scientist Development Award and a Howard Hughes Medical Institute Early Career Grant. We also acknowledge support from National Institutes of Health Grants Digestive Disease Center DK56339 (H.P.J.), P01 HG000205 (H.P.J.), NCI Innovative Molecular Analysis Technology 1R33CA1745701 (H.P.J.), NHGRI R01 HG006137 (H.P.J.). Other support came from the Gastric Cancer Foundation (H.P.J.).

## Author contributions

JC and HPJ conceived and designed the study. JC performed all the experiments. JC, BL, NA, and SG conducted the sequencing data analysis. CH was involved in the development of the organoids. JC and CWB conducted sequencing. HPJ supervised and coordinated all aspects of the analysis and experiments. All authors revised, read, and approved the final manuscript.

